# Distributed disruption and remodelling of synaptome architecture following traumatic brain injury

**DOI:** 10.64898/2025.12.19.695532

**Authors:** Aimun A.B. Jamjoom, Ragini Gokhale, Jonathan Rhodes, Zhen Qiu, Seth G.N. Grant

## Abstract

A single cortical impact in mice altered the number and molecular composition of excitatory synapses across 222 brain regions. Early effects targeted synapses with rapid protein turnover, followed by changes in slow-turnover synapses. This injury-induced remodelling of synaptome architecture mirrors responses to genetic mutation, ageing, sleep deprivation and experience, suggesting that reorganisation of excitatory synapse populations is a shared mechanism underlying brain plasticity across diverse conditions.

## MAIN TEXT

Traumatic brain injury (TBI) is a leading global cause of death and long-term disability across all age groups^1^. Our current knowledge of how TBI affects synapses remains limited to studies of small numbers of synapses in select brain regions^2–6^. The development of synaptome mapping has enabled systematic, high-resolution analysis of billions of individual synapses across the whole mouse brain, giving rise to the concept of the synaptome – the full repertoire of molecularly diverse synapse types – and its spatial and temporal configuration, referred to as synaptome architecture^2,7–12^. Synaptome mapping uses genetically tagged synaptic proteins to classify synapses into distinct types and subtypes based on molecular composition, protein turnover and nanoarchitecture. In this study, we examine the effects of TBI (lateral fluid percussion injury, LFPI) and craniotomy (sham-LFPI), a widely used method for creating cranial windows for live imaging of the brain, on synaptome architecture (Fig. 1a).

**Figure 1:**
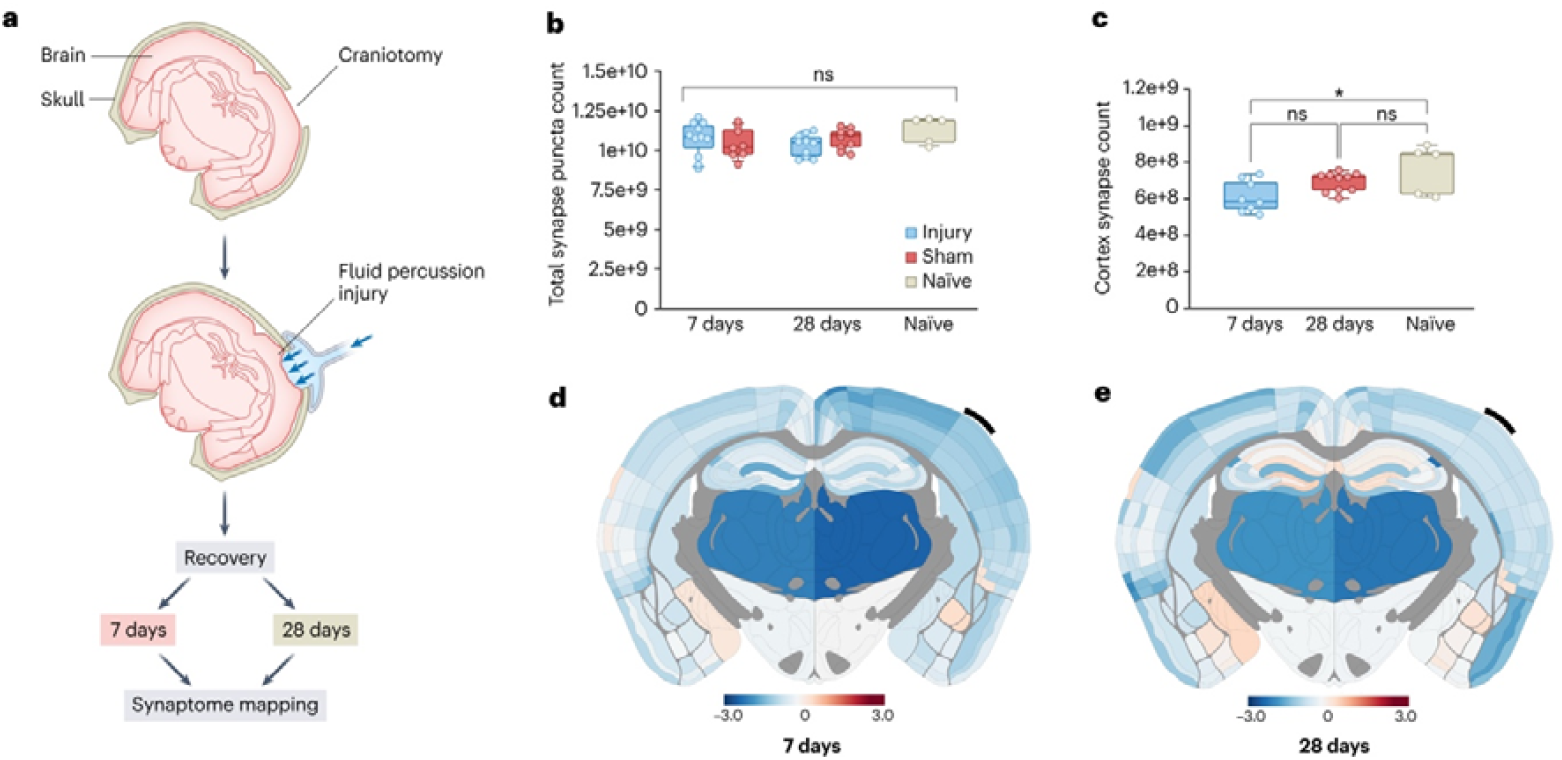
Impact of lateral fluid percussion injury on the mouse brain synaptome. **(a)** Lateral fluid percussion injury (LFPI) experimental pipeline. **(b)** Comparison of whole-brain synapse puncta count across mouse cohorts (injury, sham-LFPI and surgery-naïve) and time points (7 and 28 days) (ns, p>0.05; upper and lower lines indicate the 95th and 5th percentiles). **(c)** Comparison of synapse puncta count in ipsilateral cortex underlying craniotomy (MOp, MOs and SSp-tr) between sham-LFPI and surgery-naïve groups (*, p<0.05; one-way ANOVA with Tukey multiple comparisons test; upper and lower lines indicate the 95th and 5th percentiles), **(d**,**e)** Spatial heatmaps of effect size (Cohen’s *d*) comparing sham-LFPI and surgery-naïve groups at 7D (d) and 28D (e). Black line, location of injury site.

We applied previously validated tools and protocols for brainwide synaptome mapping of excitatory synapse populations^7–11^. Transgenic mice expressing PSD95-GFP and SAP102-mKO2 fusion proteins were used to fluorescently label postsynaptic scaffold proteins crucial for the assembly of signalling complexes at excitatory synapses^13–15^ and essential for synaptic plasticity and cognitive function^16–18^. A group of mice (N=21) received a single LFPI pulse (1.0–1.5 atm) delivered to the intact dura over the left parietal cortex; a second group (N=18), referred to as sham-LFPI, received a craniotomy which maintained the dura mater intact but without the application of the fluid percussion; and a third group (N□=□5) were surgery-naïve (Fig. 1a). The LFPI and sham-LFPI group were analysed at 7 days (7D) and 28 days (28D) post-injury. Mid-coronal brain sections at Bregma −1.9 mm were imaged using spinning disc confocal microscopy at 100× magnification (optical resolution ~260□nm; pixel size 84□nm). Synaptic puncta were quantified in each mouse across 222 anatomically defined subregions (111 subregions in the ipsilateral and contralateral hemispheres to the LFPI). The images of synaptic puncta were processed with the SYNMAP synaptome mapping analysis pipeline, which detects individual excitatory synapses and classifies them into three principal types: Type 1 expressing only PSD95, Type 2 expressing only SAP102, and Type 3 expressing both PSD95 and SAP102. These three types were further subdivided into 37 molecularly and morphologically distinct synapse subtypes based on features including protein abundance, shape and size. Of these, 30 subtypes were additionally stratified into those with short protein lifetime (SPL) and long protein lifetime (LPL) categories based on the known protein lifetime of PSD95^7^.

Across the cohorts, the median whole-brain synapse puncta count was 1.075×10^1^□ with an interquartile range (IQR) of 1.41×10□ (Q1 = 9.87×10□, Q3 = 1.13×10^1^□). There was no statistical difference in the whole-brain puncta count across the cohorts at 7D and 28D (p>0.05; one-way ANOVA with Tukey multiple comparisons test) (Fig. 1b). Next, we compared the sham-LFPI and surgery-naïve groups to ask if craniotomy alone affected the number of excitatory synapses in the brain. In the isocortex beneath the craniotomy (primary motor area, secondary motor area and primary somatosensory area), a significant reduction in total puncta count was observed at 7D (p<0.05; one-way ANOVA with Tukey multiple comparisons test), but not at 28D (Fig 1c). Brain maps showing effect sizes (Cohen’s *d*) between sham-LFPI and surgery-naïve revealed loss of synapse density within the thalamus at both 7D and 28D post-injury, indicating the effect of craniotomy extends to subcortical regions (Fig. 1d,e). These results show that craniotomy alone, even when the dura is left intact, can lead to loss of synapses in local and distant brain regions.

We next examined LFPI in comparison to the surgery-naïve group and, as expected, found that at 7D the regions with greatest synapse loss were in the isocortex underlying the injury site (p<0.05, Bayesian estimation with Benjamini–Hochberg correction) (Fig. S1). By 28D there was widespread reduction in puncta density across the isocortex and hippocampus in both hemispheres. Comparison of LFPI and sham-LFPI groups showed little difference at 7D (p>0.05, Bayesian estimation with Benjamini–Hochberg correction) (Fig. 2a and Fig. S2). However, this difference increased by 28D, with LFPI causing a significant decrease in the density of synapses expressing PSD95 in the contralateral isocortex and the hippocampus (p<0.05, Bayesian estimation with Benjamini-Hochberg correction) (Fig. 2b,c).

**Figure 2:**
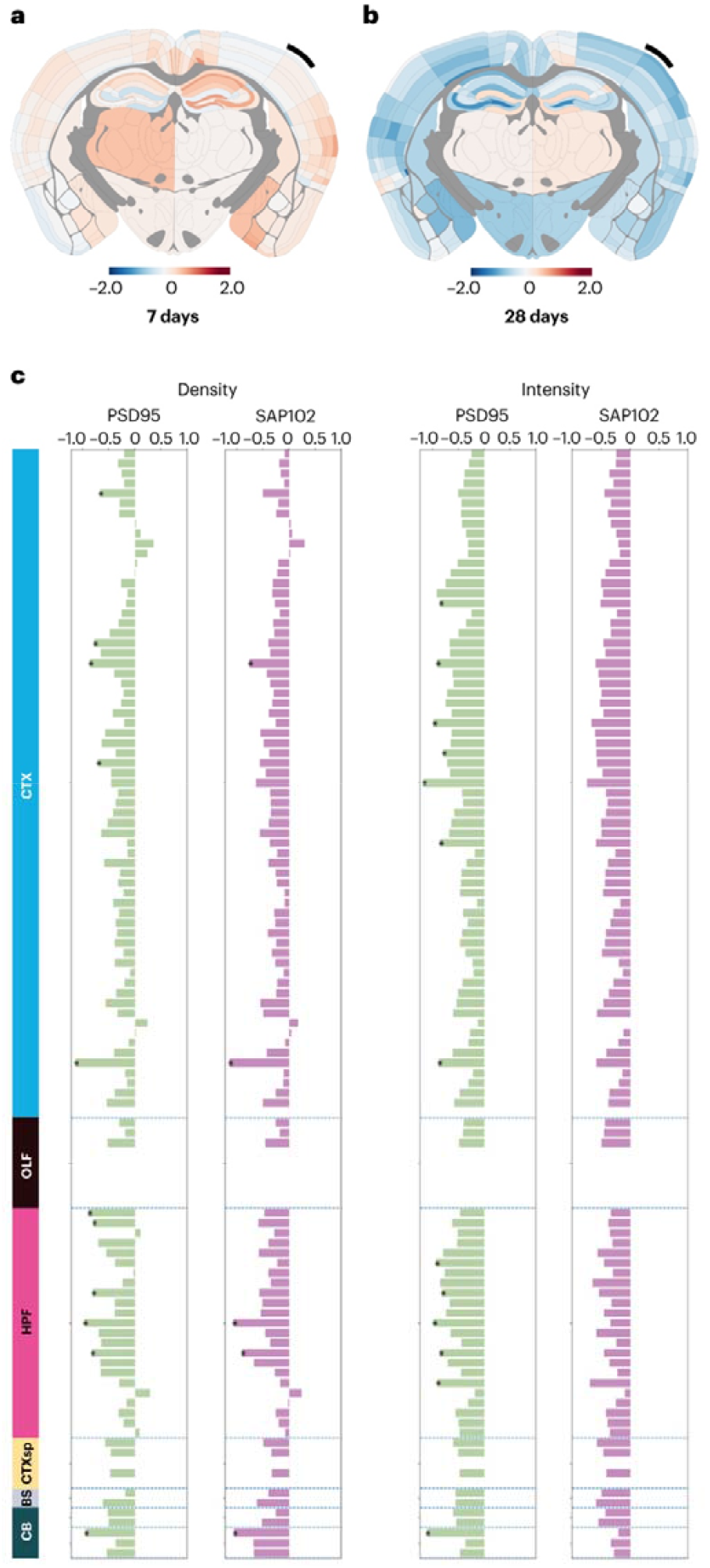
LFPI leads to delayed disruption of the synaptome that spreads to the contralateral hemisphere. **(a**,**b)** Spatial heatmaps of effect size (Cohen’s *d*) comparing LFPI (injury) and sham-LFPI groups at (a) 7D and (b) 28D. Black bars, location of craniotomy. **(c)** Cohen’s d effect size difference between LFPI and sham-LFPI in puncta density and intensity in the contralateral hemisphere at 28D (*p<0.05, Bayesian estimation with Benjamini– Hochberg correction). CTX: Cerebral cortex; OLF: Olfactory Area; HPF: Hippocampal Formation; CTXsp: Cerebral cortex, splenial region; BS: Brainstem; CB: Cerebellum

In addition to measuring the density of PSD95 and SAP102 expressing synapses, we measured their protein content (intensity) and size. LFPI resulted in changes in the mean intensity and size of excitatory synapse populations in each brain region (Fig. S2). To explore these changes in the synapse populations in greater detail, we created heatmaps showing the effect size (Cohen’s *d*) of LFPI compared with sham-injury on the density for each synapse type and subtype in 111 subregions in each hemisphere at 7D and 28D (Fig. S3). These heatmaps reveal LFPI causes particular subtypes to increase and others to decrease in density in a highly distributed manner at both 7D and 28D. These anatomically widespread and synapse-specific changes are informative regarding the brain’s response to injury. For example, Type 1 subtypes 2-5 are decreased in cortical areas near the injury site at 7D, and by 28D show widespread decrease across all brain regions in both hemispheres. The Type 2 subtypes 12-15 are decreased in ipsilateral and contralateral cortices at 7D and 28D, suggesting a particular vulnerability. The Type 3 subtypes 20-31 are reduced at 7D but reverse to increase at 28D in the ipsilateral cortex. Within any given heatmap there are increased and decreased subtypes, suggesting that the molecular identity of the decreased subtypes has switched to become that of the subtypes that increase. These results show that brain injury induces local and distributed changes in the density of synapse types and subtypes which evolve with time after the injury.

The dynamic evolution of synaptome architecture that we observed following brain injury may inform mechanisms of synaptic resilience and repair. We hypothesised that the changes in synapse molecular composition occurring between 7D and 28D would require protein turnover, and thus synapses with different rates of protein turnover might show differential responses during this period. To examine the relationship between the temporal evolution of the injury response and changes in the density of subtypes with different rates of PSD95 turnover^7^, we first ranked the 37 subtypes according to the significance of their change (increase or decrease) (p<0.05, Bayesian estimation with Benjamini–Hochberg correction) and marked the top 6 SPL (6, 8, 11, 28, 29, 31) and top 6 LPL (2, 3, 5, 20, 30, 34) subtypes (Fig. 3a).

**Figure 3:**
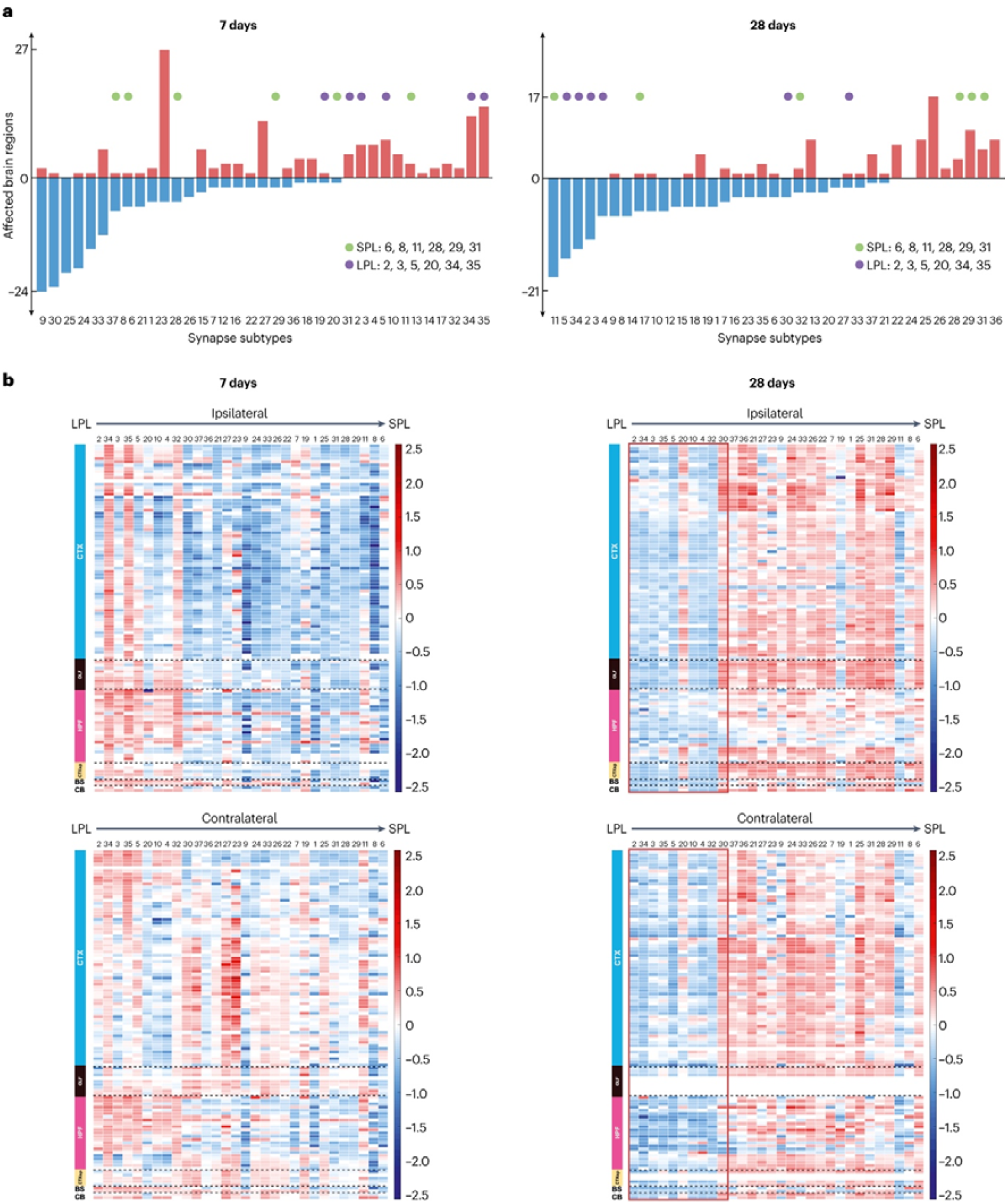
LFPI differentially impacts synapse subtypes with different protein lifetimes. Comparison between LFPI and sham-LFPI groups showing **(a)** Ranking of synapse subtypes according to the number of brain subregions significantly affected in injury mice at 7D and 28D (p<0.05, Bayesian test with Benjamini–Hochberg correction). **(b)** Heatmap (Cohen’s *d* values, key on right) of brain regions and synapse subtypes comparing LFPI and sham-LFPI groups ranked from left to right according to PSD95 protein lifetime (LPL, longest protein lifetime on left and SPL, shortest protein lifetime on right) across 111 brain subregions (both ipsilateral and contralateral) across two time points (7 and 28 days). LPL synapses are boxed in red in 28D samples.

Next, heatmaps were plotted, ranked according to protein lifetime (Fig. 3b). These plots show several striking features. As shown in the heatmaps in Figure 3b, at 28D there is a blue vertical stripe (boxed in red) encompassing the leftmost 10 subtypes in both ipsilateral and contralateral hemispheres, indicating that by 28D there is widespread reduction in LPL synapses. Correspondingly, there is a general increase in the number of SPL synapse subtypes. Moreover, this pattern is not present at 7D, indicating that it evolves after the first week following injury. Indeed, at 7D there is a reduced density of SPL synapses, particularly within the ipsilateral isocortex, indicating that these synapse subtypes underpin the synapse density loss described above. Collectively, this analysis highlights an initial decrease in SPL synapses and increase in LPL synapses at 7D and the reversal of this relationship by 28D.

Our study reveals progressive and widespread reorganisation in the synaptome architecture that extends from the initial impact site across the brain over a 28-day period. We uncovered selective vulnerability of synapse types and subtypes that were region and time specific. Moreover, when examining synapse protein lifetime, a distinct two-phase profile emerged. At 7 days there was early loss of SPL synapses combined with an increase in LPL synapses, and in the delayed phase at 28 days this relationship was reversed. This initial impact and subsequent shift reflects an underlying differential vulnerability to injury of SPL and LPL synapses and the role of proteostasis in reorganising the proteomes of individual synapses to remodel the overall population. Such changes may underpin sequelae of brain injury including memory loss and alterations in proteostasis linked to neurodegeneration. Together with upregulation of synaptic gene expression after head trauma^19^, changes in synaptic proteostasis may result in the observed changes in synapse proteome composition and synaptome architecture.

The present study further reveals that mechanisms of synaptic vulnerability and repair following traumatic brain injury share common features with adaptive synaptome responses to mutation, ageing, sleep deprivation, and experience^7,9,11,12^. Our previous synaptome analyses across these conditions have identified conserved organisational principles of excitatory synapse populations: synapses with specific molecular compositions are selectively targeted; adaptive responses involve a rebalancing of SPL and LPL synapses; and these changes are expressed brain-wide or across extensive anatomical territories. The conservation of these mechanisms implies that synaptome responses to TBI are modulated by age, sleep, and experience. Consistent with this prediction, increasing age is a well-established risk factor for poor outcome^20,21^, environmental enrichment promotes rehabilitation^22^, and sleep disturbances adversely affect the clinical course of TBI recovery^23,24^. Our study also provides insights into how the cognitive and behavioural impairments arise from TBI since the synaptome architecture is a mechanism for encoding innate and learned behavioural representations^7,10,12,25,26^ and its disruption and reorganisation would result in altered cognition and behavioural phenotypes. Together, these findings position synaptome mapping as a powerful framework for elucidating the mechanisms of selective synaptic vulnerability and as a platform for the development of diverse, mechanism-informed therapeutic interventions.

## METHODS

### Animals

All animal experimentation was undertaken in accordance with Home Office guidelines [Animal (Scientific Procedures) Act 1986] and was approved by The University of Edinburgh. PSD95-eGFP and SAP102-mKO2 knock-in mice were used. The gene targeting and mouse generation were described previously^10^. Males homozygous for PSD95-eGFP and hemizygous for SAP102-mKO2 and aged 8-16 weeks were included in the experiments. The mice were allowed to acclimatize in the animal unit for at least 7 days prior to experimentation. The mice were kept in a 12-hour light/dark cycle and given free access to food and water before and after the experimental procedure.

### Lateral fluid percussion injury model

The mice were anaesthetized with 5% isoflurane (Merial, UK) in 100% O_2_ for 2 minutes prior to surgery. The mice were then weighed and placed in a stereotaxic frame (Model 900, David Kopf Instruments). A nose cone was used to maintain anaesthesia. During surgery, mouse body temperature was maintained at 37±0.5°C using a heat pad and rectal thermometer probe (Homoeothermic Blanket Control Unit, Harvard Apparatus). A razor was then used to shave a small patch of hair from the top of the mouse’s head and eye ointment (Xailin) was applied to the eyes. A midline scalp incision was fashioned and the skin flaps reflected using skin clips. A plastic guide was glued onto the skull using 3M Vetbond and a lateral 2 mm circular craniectomy was then trephined (Miltex) midway between the bregma and lambda. Care was taken not to injure underlying dura. An injury hub was created by cutting off the female end of a Luer-Loc needle hub. This was secured over the craniectomy site using glue (Loctite Power Flex Gel) and dental cement (Simplex Red, Associated Dental Products). The mice were then removed from the stereotactic frame and the depth of anaesthesia reduced until they responded to tail pinch. The mice were then placed in the anaesthetic box for 1 minute. Once complete, the injury hub was filled with normal saline and attached to the fluid percussion device (Custom Design and Fabrication, Virginia Commonwealth University). When they showed response to tail stimulus (to standardize anaesthesia levels), a mild to moderate injury (1.1-1.5 atm) was inflicted. This was determined by oscilloscope attached to the fluid percussion device. Sham mice had the exact same procedure except the release of the pendulum to inflict the percussion injury was omitted.

Following this, the mice were disconnected from the fluid percussion device and placed on their back to measure their righting time - a marker of the neurological severity of the injury. Exclusion criteria were defined as follows: dural tear during craniectomy; dural rupture following application of fluid percussion or post-injury paralysis lasting greater than 30 minutes. Following righting, the mice were then re-anesthetized and had the hub removed and the skin incision sutured using 5-0 Vicryl (Ethicon). A 2 mg/kg dose of Bupivacaine (AstraZeneca) was applied to the wound to manage pain. Mice were then placed in a heated recovery chamber at 30°C (Lyon Electric Company) for 30 minutes prior to being returned to a cage. Mice were followed up to 7 days (7D) or 28 days (28D). Mice were initially monitored daily for the 5 days postoperatively. Following this, they were assessed once a week until the end of the experiment.

### Tissue collection and processing

Mice were administered a terminal dose of 0.1 ml 20% phenobarbital sodium (Euthatal – Merial Animal Health) by intraperitoneal injection. Once the mice were fully anaesthetized (not responding to tail pinch), they were pinned to a cork board and underwent a thoracotomy. The heart was fully exposed and a small knick was made in the right atrium. Subsequently, 10 ml phosphate buffered solution (PBS, Fisher Scientific) was injected slowly into the left ventricle. This was followed by 10 ml 4% paraformaldehyde (PFA, prepared by diluting 16% Alfa Aesar 1:4 in PBS). The mice were then decapitated and the brain dissected free from the skull. It was post-fixed in 5 ml 4% PFA for 3-4 hours and then placed in a 30% sucrose solution (prepared by diluting sucrose (VWR Chemicals) in PBS) for 48 hours. The brains were then placed in a solution of 50:50 30% sucrose and Optimum Cutting Temperature medium (OCT, VWR International) for 1 hour before being embedded in a mould (Sigma-Aldrich) in OCT. This was achieved by immersing the moulds in a beaker of isopentane (Sigma-Aldrich) which was then placed in liquid nitrogen. Once frozen, the brains were kept at −80°C until sectioned. A NX70 Cryostat (Thermo Scientific) was used to cut coronal sections 18 microns in depth. A small drop of PBS was placed on a Superfrost Plus glass slide (Thermo Scientific) before sections were collected. Slides were left to dry at 4°C overnight, then stored at −80°c until further analysis. Sections were kept away from light to minimize loss of fluorescence.

### Spinning disc confocal microscopy

Image acquisition was performed using the Andor Revolution XDi system using a Dual Nipkow disk (CSU-W1) which combines two pinholes of 25 and 50 μm. We utilized two laser wavelengths: 488 nm (eGFP) and 561 nm (mKO2). Light from the lasers was directed onto the sample using a dichroic mirror and quad filter. A 2x post-magnification lens was placed in front of the camera to increase sampling rate. The system also contained the Borealis Perfect Illumination Delivery which improves the uniformity of the field illumination. Images were acquired using an Olympus UplanSAPO 100x oil-immersion lens (N.A 1.4) and detection was achieved using the Andor iXon Ultra monochrome back-illuminated EMCCD. Acquired images were 512 x 512 pixels and 16-bit depth. For whole brain acquisition, images underwent mosaic tilling across a grid defined by the user. Within this grid, focus points were placed by the user which allowed for adaptive focusing throughout the imaged sample. No z-stacking was taken in images.

### Synaptome mapping pipeline

The synaptome mapping (SYNMAP) pipeline^8,10^ was developed and standardized to systematically map individual PSD95-eGFP and SAP102-mKO2 puncta across the entire brain. Using in-house developed deep-learning methods^27,28^, SYNMAP comprises a sequence of automated image analysis procedures that includes puncta detection, colocalization, classification and map reconstruction. To define the anatomical regions, manual delineation was employed using the Allen Mouse Brain Atlas as a guiding resource. For each punctum, intensity, size and morphological features were extracted at single-synapse resolution. These features were then used as inputs to an in-house unsupervised model that generated a hierarchical synapse catalogue comprising three primary classes (PSD95 only, SAP102 only, colocalized) and 37 subtypes. In our previous study^7^, we quantified PSD95 protein lifetimes across 30 synapse subtypes that express PSD95 by tracking the temporal persistence of labelled PSD95 molecules. The six subtypes with the longest turnover times were classified as long protein lifetime (LPL) synapses, and the six with the shortest turnover times were classified as short protein lifetime (SPL) synapses.

### Statistical analysis

#### Cohen’s d formula

Cohen’s d effect size was calculated as per Cohen^29^ as follows:

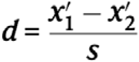

where is the mean of one of the groups and is the pooled standard deviation as:

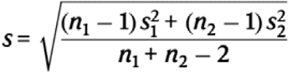

where is the pooled standard deviation as:

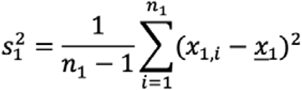

#### Bayesian analysis

Bayesian estimation^30^ was used to provide an objective assessment of the effects of traumatic injury on synaptome features, including PSD95 parameters, subtype densities and diversity. The results were corrected over all subregions using the Benjamini-Hochberg procedure.

## ACKNOWLEDGEMENTS

We extend our appreciation to K. Elsegood and C. McLaughlin for laboratory support; D. Dominic for assistance in data management; C. Davey for editing; and D. Maizels for artwork. Funding was provided by a Wellcome Trust Translational Medicine and Therapeutics grant (106851/Z/15/Z) to A.A.B.J.

## AUTHORS CONTRIBUTIONS

A.A.B.J. conceived the study, developed the methodology, performed validation, formal analysis, and investigation, and led project administration and funding acquisition. A.A.B.J. drafted the original manuscript and contributed to manuscript review, editing, and data visualization. R.G. and Z.Q. contributed to methodology development, software implementation, formal analysis, data curation, and data visualization, and participated in manuscript review and editing. J.R. contributed to methodology development and provided resources and reviewed and edited the manuscript. S.G.N.G. contributed to study conceptualization and methodology, provided resources, supervised the project, acquired funding, and contributed to manuscript drafting, review, and editing.

## SUPPLEMENTARY FIGURES

**Figure S1:**
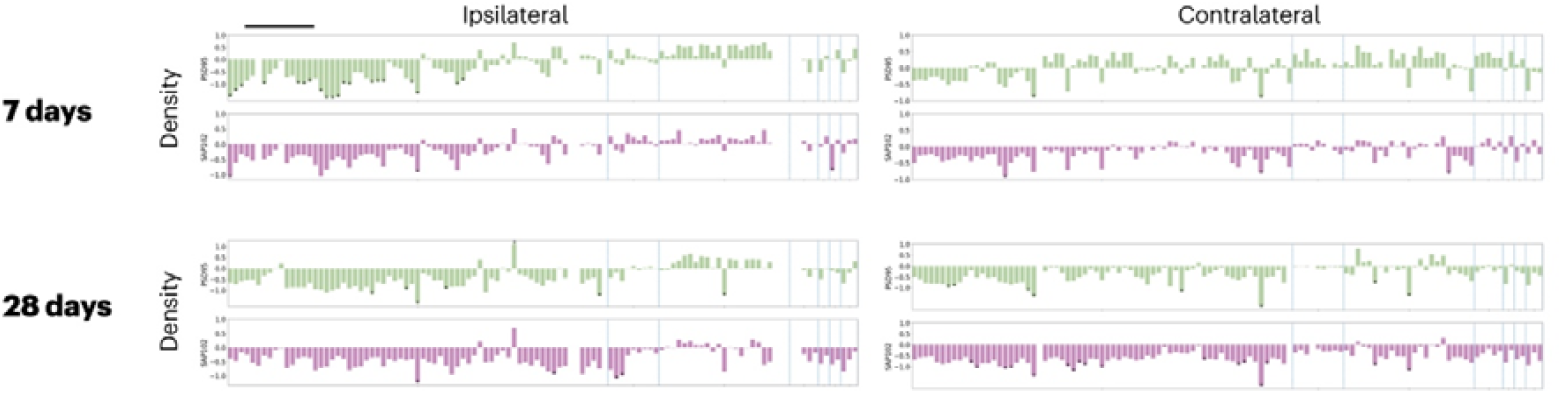
Cohen’s d effect size of puncta density between injury (LFPI) and surgery-naïve groups at 7D and 28D (*p<0.05, Bayesian test with Benjamini– Hochberg correction). Black line, location of injury site.

**Figure S2:**
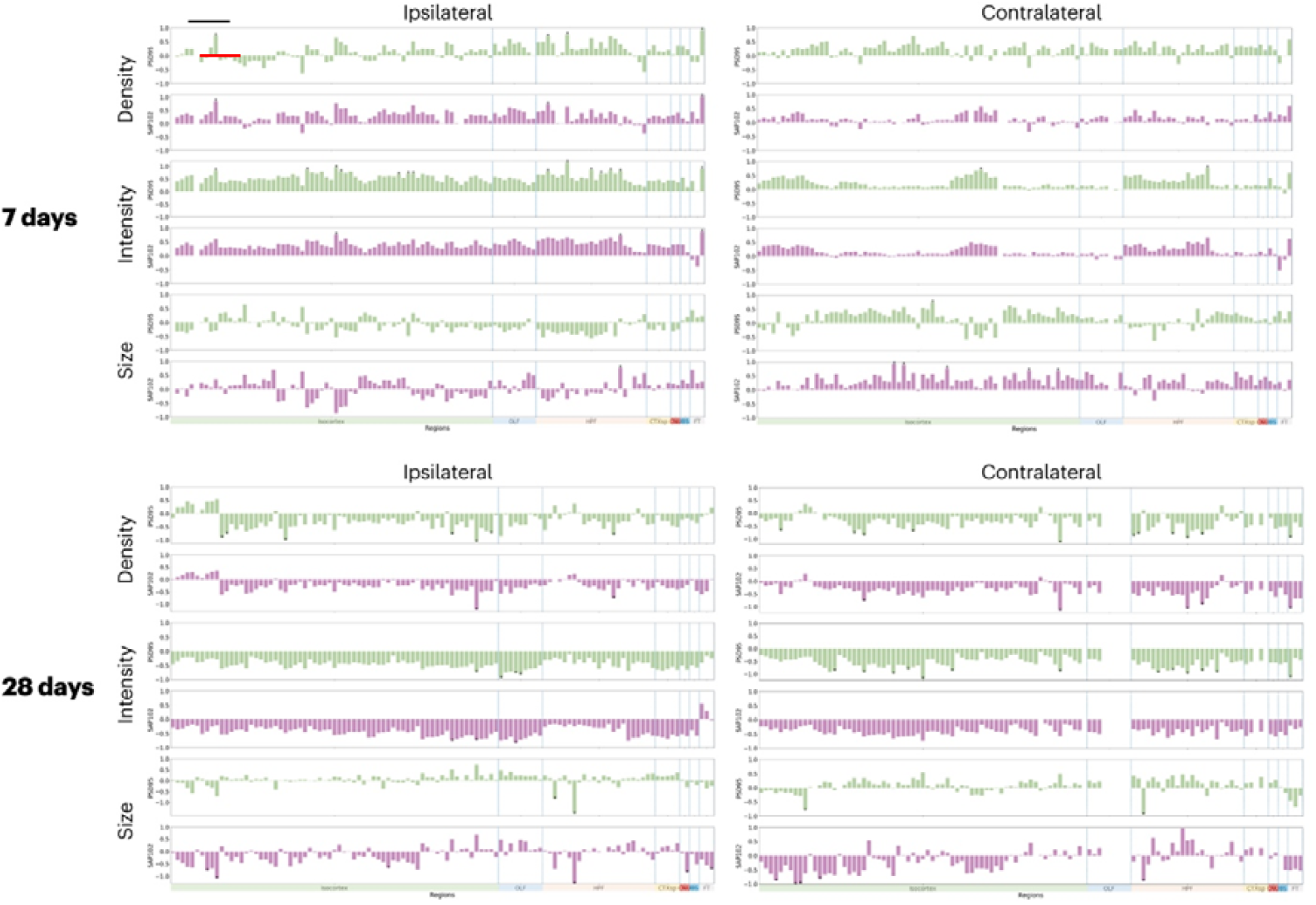
Cohen’s d effect size of puncta density, intensity and size between injury (LFPI) and sham-LFPI groups at 7D and 28D (*p<0.05, Bayesian test with Benjamini–Hochberg correction). Black line, location of injury site.

**Figure S3:**
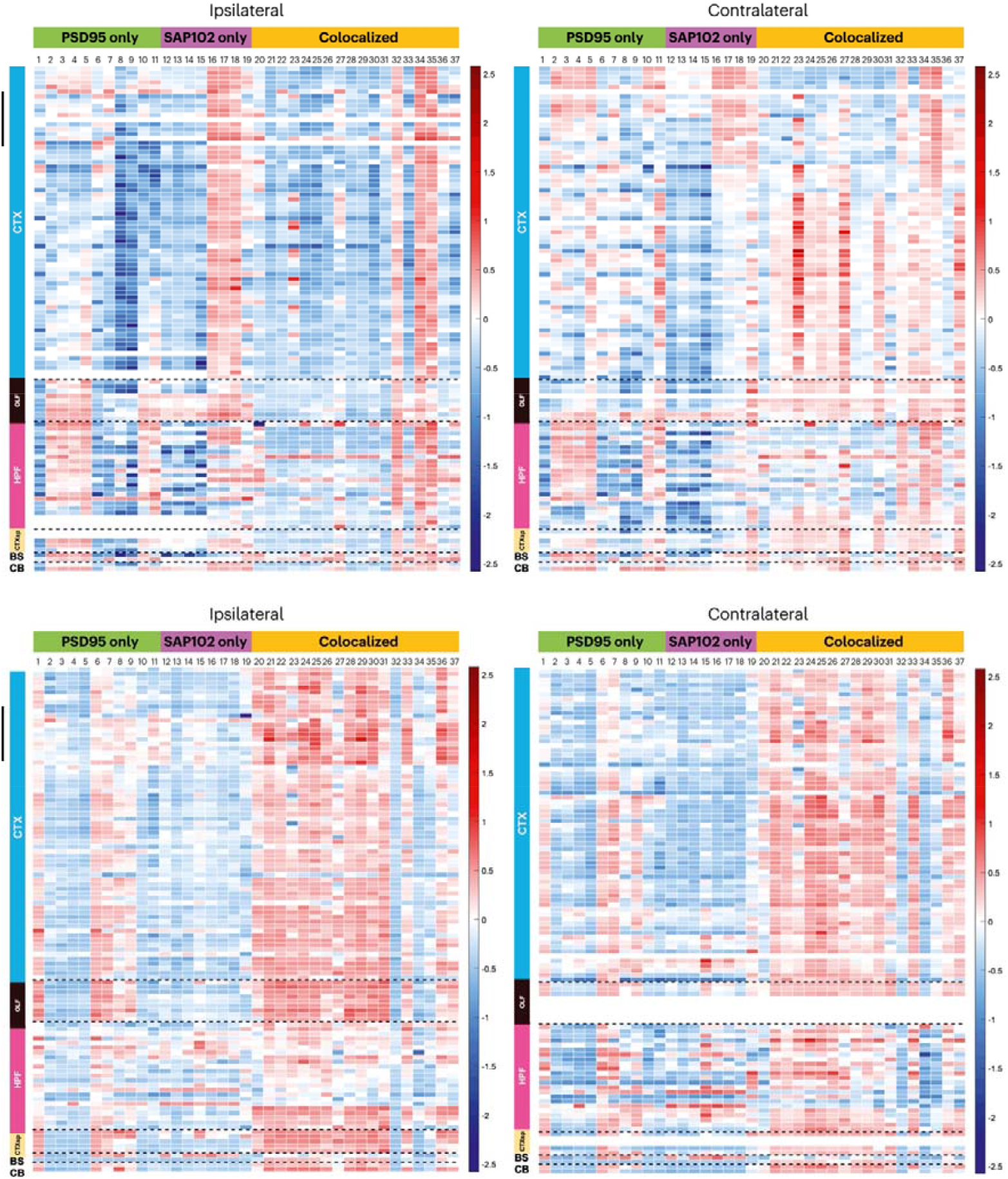
Heatmap of Cohen’s d effect size (key on right of each plot) of puncta density between injury (LFPI) and sham-LFPI groups at 7D and 28D categorised by synapse type and subtype. Black line, location of injury site.

## Notes

### Competing Interest Statement

The authors have declared no competing interest.

